# A common flanking variant is associated with enhanced meiotic stability of the *FGF14*-SCA27B locus

**DOI:** 10.1101/2023.05.11.540430

**Authors:** David Pellerin, Giulia Del Gobbo, Madeline Couse, Egor Dolzhenko, Marie-Josée Dicaire, Adriana Rebelo, Virginie Roth, Marion Wandzel, Céline Bonnet, Catherine Ashton, Phillipa J. Lamont, Nigel G. Laing, Mathilde Renaud, Gianina Ravenscroft, Henry Houlden, Matthis Synofzik, Michael A. Eberle, Kym M. Boycott, Tomi Pastinen, All of Us Long Reads Working Group, Bernard Brais, Stephan Zuchner, Matt C. Danzi

**Affiliations:** Department of Neurology and Neurosurgery, Montreal Neurological Hospital and Institute, McGill University, Montreal, QC, Canada; Department of Neuromuscular Diseases, UCL Queen Square Institute of Neurology and The National Hospital for Neurology and Neurosurgery, University College London, London, United Kingdom; Children’s Hospital of Eastern Ontario Research Institute, University of Ottawa, Ottawa, ON, Canada; Centre for Computational Medicine, The Hospital for Sick Children, University of Toronto, Toronto, ON, Canada; Pacific Biosciences, Menlo Park, CA, USA; Dr. John T. Macdonald Foundation Department of Human Genetics and John P. Hussman Institute for Human Genomics, University of Miami Miller School of Medicine, Miami, FL, USA; Laboratoire de Génétique, CHRU de Nancy, France; INSERM-U1256 NGERE, Université de Lorraine, Nancy, France; Department of Neurology, Royal Perth Hospital, Perth, WA, Australia; Centre for Medical Research University of Western Australia and Harry Perkins Institute of Medical Research, Perth, Western Australia, Australia; Service de Neurologie, CHRU de Nancy, France; Service de Génétique Clinique, CHRU de Nancy, France; Division of Translational Genomics of Neurodegenerative Diseases, Hertie-Institute for Clinical Brain Research and Center of Neurology, University of Tübingen, Tübingen, Germany; German Center for Neurodegenerative Diseases (DZNE), Tübingen, Germany; Genomic Medicine Center, Children’s Mercy Kansas City, Kansas City, MO, USA; UMKC School of Medicine, University of Missouri Kansas City, Kansas City, MO, USA; Department of Human Genetics, McGill University, Montreal, QC, Canada

**Keywords:** *FGF14*, spinocerebellar ataxia 27B, ataxia, spinocerebellar ataxia, GAA repeat instability, trinucleotide repeats, meiotic instability, repeat expansion disorder

## Abstract

The factors driving initiation of pathological expansion of tandem repeats remain largely unknown. Here, we assessed the *FGF14*-SCA27B (GAA)•(TTC) repeat locus in 2,530 individuals by long-read and Sanger sequencing and identified a 5’-flanking 17-bp deletion-insertion in 70.34% of alleles (3,463/4,923). This common sequence variation was present nearly exclusively on alleles with fewer than 30 GAA-pure repeats and was associated with enhanced meiotic stability of the repeat locus.

## Main Text

Dominantly inherited (GAA)•(TTC) repeat expansions in intron 1 of the fibroblast growth factor 14 (*FGF14*) gene have recently been shown to cause spinocerebellar ataxia 27B (SCA27B; MIM: 620174)^1,2^. Initial observations suggest that expanded alleles are unstable during intergenerational transmission^1^, although the underlying mechanisms driving this instability remain unknown. Here, we provide large-scale evidence that a common sequence variation flanking the *FGF14* repeat locus is present nearly exclusively on short GAA-pure alleles and is associated with enhanced meiotic stability of the repeat locus.

We first analyzed the repeat length, motif content, and flanking sequences of the *FGF14* repeat locus in a set of 541 alleles from 339 individuals of self-reported European ancestry (146 controls and 193 patients with late-onset ataxia, including 32 patients with SCA27B for whom results were independently validated by nanopore sequencing) using Sanger sequencing (see Online Methods). We defined the flanking sequences as the 25 nucleotides immediately adjacent to the *FGF14* repeat locus (GRCh38, chr13:102,161,575-102,161,726) on both 5’ and 3’ ends. We found that only 2 of the 541 alleles (0.37%) perfectly matched the reference 5’-flanking sequence, while 186 additional alleles (34.38%) were within 3 single nucleotide variations (substitution, insertion, or deletion) of the 25-bp reference 5’-flanking sequence. Though we will hereafter refer to both the 25-bp reference 5’-flanking sequence and its variants carrying no more than 3 single nucleotide variations as the reference 5’-flanking sequence (5’-RFS), it is not the major allele at the *FGF14* repeat locus. The majority of the 541 alleles (343, 63.40%) contained a 17-bp deletion-insertion (NC_000013.11: g.102161566_102161576delinsTAGTCATAGTACCCCAA) in the 5’-flanking sequence of the repeat locus. Only ten alleles (1.85%) had 5’-flanking sequences that did not contain the 17-bp deletion-insertion and differed by more than 3 single nucleotides from the 25-bp reference 5’-flanking sequence (Figure 1A and 1B). Remarkably, the 17-bp deletion-insertion common 5’-flanking variant (5’-CFV) was exclusively observed in alleles carrying fewer than 30 GAA-pure repeats whereas the 5’-RFS and other 5’-flanking sequences were only present in alleles with more than 30 repeats, of which only 31.81% (63/198) were GAA-pure (Figure 1C). Alleles shorter than 30 repeats were perfectly separated from longer alleles by the presence of the 5’-CFV, such that none of the expanded, pathogenic (GAA)_≥250_ alleles carried the 5’-CFV.

**Figure 1:**
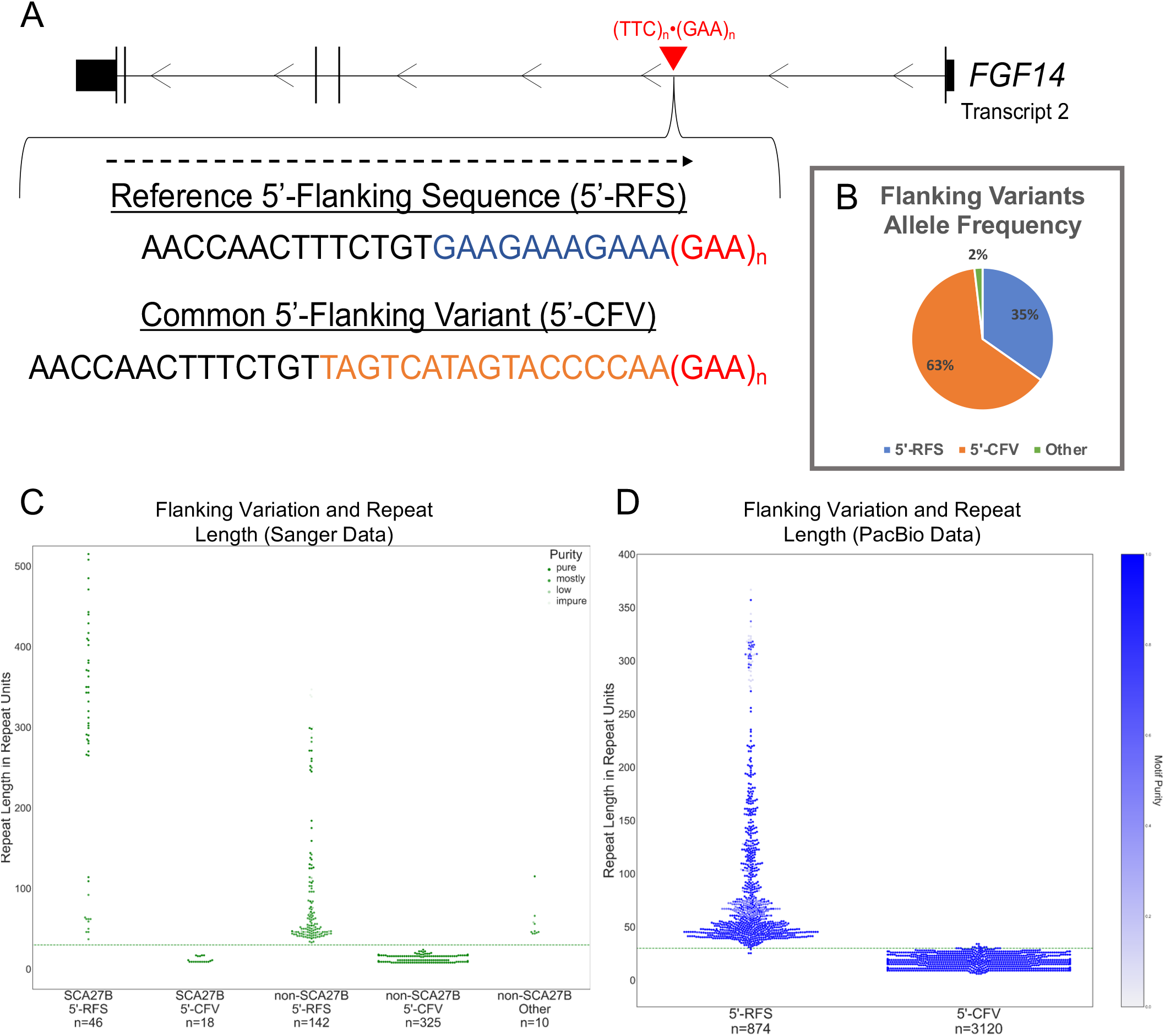
A common 5’-flanking sequence variant is associated with smaller *FGF14* GAA repeat size. A) Diagram of the *FGF14* gene with the location of the (GAA)_n_•(TTC)_n_ repeat locus in the first intron. The reference 5’-flanking sequence (5’-RFS) and the 17-bp deletion-insertion common 5’-flanking variant (5’-CFV) are shown. The portion of the 5’-RFS deleted in the 5’-CFV is shown in blue. The 5’-CFV consists of a 17-bp deletion-insertion (NC_000013.11: g.102161566_102161576delinsTAGTCATAGTACCCCAA), shown in orange. The 5’-flanking sequence is presented relative to the positive strand for clarity. B) Pie chart of allele frequency of the 5’-RFS, 5’-CFV, and other flanking variants in 541 alleles analyzed by Sanger sequencing. C) Swarmplot of GAA repeat size as estimated by Sanger sequencing for 541 alleles shows that the 5’-CFV is consistently associated with alleles containing fewer than 30 GAA repeats while the 5’-RFS is associated with larger alleles, including pathogenic ones. Each of the two alleles of patients with SCA27B are shown in either the 5’-RFS or 5’-CFV categories, even though only (GAA)_≥250_ alleles are pathogenic (all carrying the 5’-RFS). The color of the data points is a function of the GAA repeat motif purity, with dark green indicating pure and lighter green impure motif (a hue scale is shown on the top right corner of the graph). Motif purity of the repeat locus was assessed using the following ordinal scale: impure (non-GAA motif), low purity (<75% GAA motif), mostly pure (75-99% GAA motif), and pure (100% GAA motif). D) Swarmplot of GAA repeat size as estimated by PacBio HiFi sequencing for 3,995 alleles shows a similar pattern. Alleles possessing any other flanking sequences and two alleles of over 800 repeat units carrying the 5’-RFS were excluded (see Supplementary Figure 1). The color of the data points is a function of the GAA repeat motif purity, with dark blue indicating pure and lighter blue impure motif (a hue scale is shown on the right y axis).

We next sought to replicate these findings using whole-genome PacBio HiFi sequencing in 2,191 individuals (4,382 alleles) spanning a wide range of genetic ancestries. See the Online Methods for a description of this cohort. We found that 874 alleles (19.95%) carried the 5’-RFS, of which only 6 alleles (0.14%) perfectly matched the 25-bp reference 5’-flanking sequence. The 17-bp deletion-insertion 5’-CFV was observed in 3,120 alleles (71.20%). As in the Sanger data, the 5’-CFV was consistently present in alleles with smaller repeat lengths than the alleles containing the 5’-RFS. Specifically, 3,103 (99.46%) of the 3,120 alleles containing the 5’-CFV had fewer than 30 GAA repeats and 3,120 (100.00%) of them were GAA-pure, while 871 (99.66%) of the 874 alleles containing the 5’-RFS had more than 30 triplets and 652 (74.60%) of them were GAA-pure (Figure 1D). These data show that the 5’-CFV is strongly associated with alleles carrying fewer than 30 GAA repeats compared to the 5’-RFS (odds ratio, 44,158; 95% confidence interval, 13,981 to 139,474; Chi-square test, *p*=0). Finally, the 5’-flanking sequence in the remaining 388 alleles (8.85%) not carrying the 5’-CFV nor the 5’-RFS displayed a range of variations that form another 6 groups (see Supplementary Results).

In a further analysis, we observed no relationship between variants in the 3’-flanking sequence of the *FGF14* repeat locus and the GAA repeat length, suggesting that only variants in the 5’-flanking region impact the *FGF14* repeat locus stability (Supplementary Results and Supplementary Figure 2). Since Friedreich ataxia is the only other known disease caused by intronic (GAA)•(TTC) repeat expansions, which are thought to arise from long normal alleles containing 12 or more triplets, but not from shorter alleles^3^, we analyzed the flanking sequence surrounding the *FXN* repeat locus in 1,027 individuals (2,054 alleles) for evidence of variants that distinguish distributions of repeat lengths. While we observed a variety of 5’- and 3’-flanking variants ranging from 1 to 11 nucleotide changes from the reference sequence, no clear segregation of allele sizes by flanking variants was observed (Supplementary Figure 3A-B).

**Figure 2:**
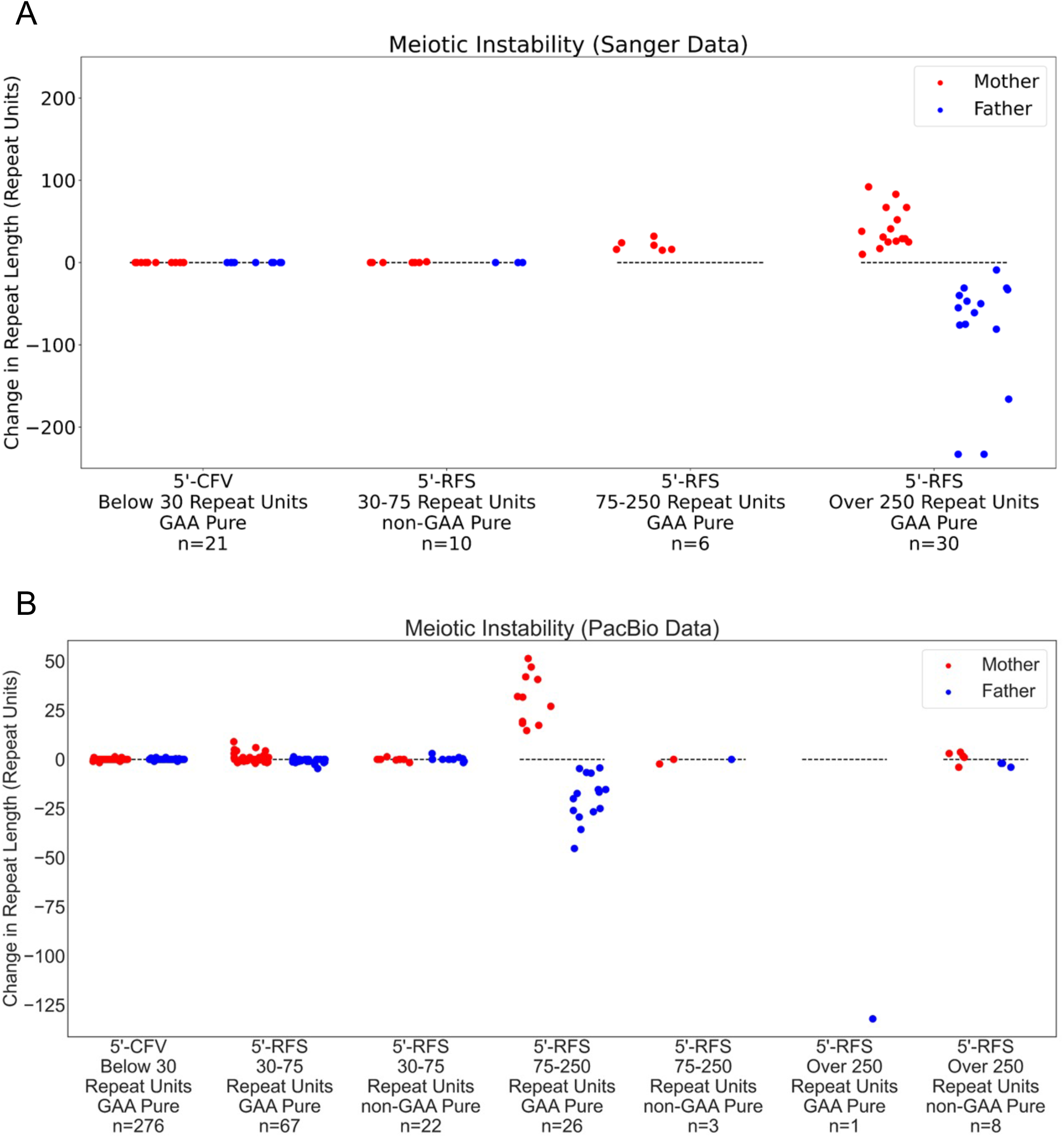
The common 5’-flanking variant enhances meiotic stability of the *FGF14* repeat locus. Strip plots of meiotic events grouped by parental allele size, 5’-flanking sequence group (5’-RFS or 5’-CFV), and GAA repeat purity as estimated by A) Sanger sequencing for 67 meioses and B) PacBio sequencing for 403 meioses. The y-axis shows the change in repeat length from parent to child. Contractions are plotted below the dashed lines while expansions are plotted above them. Random noise was applied across the x-axis within each category to allow visualization of as many data points as possible. Red dots are alleles transmitted from mother to child, while blue dots represent alleles transmitted from father to child.

We next studied the intergenerational transmission of the *FGF14* repeat locus in a total of 470 meiosis events by Sanger sequencing (67 events) and PacBio sequencing (403 events) (Figure 2A,B, Supplementary Results, and Supplementary Figures 4A,B and 5A-F). In this analysis, all alleles with fewer than 30 repeats carried the 17-bp deletion-insertion 5’-CFV, while larger alleles carried the 5’-RFS. We observed that 283 of 297 alleles (95.29%) with the 5’-CFV were stably passed from parent to offspring (Figure 2A,B). All of these 297 alleles were GAA-pure. Of the 14 non-stably transmitted alleles with the 5’-CFV, 12 differed in size by a single triplet in the offspring compared to the parent and 2 differed in size by two triplets. We did not observe a single meiotic event involving deletion of part or all of the 5’-CFV upon transmission. On the other hand, we found that alleles lacking the 5’-CFV exhibited an increasing degree of instability upon intergenerational transmission proportional to their length (Figure 2A,B). Alleles with 30 to 75 triplets and containing the 5’-RFS were stably transmitted in 50 of 99 alleles (50.51%). In this size range, 40.30% (27/67) of GAA-pure repeats were stably transmitted compared to 71.88% (23/32) of non-GAA-pure alleles. Larger alleles with 75 to 250 triplets and containing the 5’-RFS were stably transmitted in only 2 of 35 alleles (5.71%), both of which were non-GAA-pure. Finally, none of the 39 alleles larger than 250 repeats with the 5’-RFS were passed stably (Figure 2A,B). The greatest degree of instability was observed in GAA-pure alleles, with expansions observed in the female germline and contractions in the male germline, similar to what has previous been observed in Friedreich ataxia^4^. Together, these data show that the 17-bp deletion-insertion 5’-CFV is associated with greater meiotic stability than the 5’-RFS (odds ratio, 47.04; 95% confidence interval, 25.24 to 85.15; Chi-square test, *p*=2.44×10^−51^), which may account for the smaller repeat sizes associated with this flanking sequence. Our data also extend and confirm previous findings showing that GAA repeat length and purity are major factors contributing to meiotic instability^5^.

In summary, the current reference 5’-flanking sequence is in fact a minor allele at the *FGF14* repeat locus and appears to confer a risk of developing SCA27B as it is associated with enhanced meiotic instability of the repeat locus. Conversely, the 5’-CFV appears to stabilize the repeat locus and represents the major allele despite being absent from the GRCh38.p14 and T2T-CHM13v2.0 assemblies. Our data showing near-perfect separation of short (<30 GAA repeats) from long (>30 GAA repeats) alleles by the presence of the 5’-CFV suggest that deletion of this variant is likely to lead to further expansion, rather than its deletion resulting from expansion of the repeat locus. The 5’-CFV may be aiding meiotic stability via a hitherto unknown molecular mechanism, potentially linked to the formation of a DNA secondary structure that is less prone to expansion and contraction during DNA replication and transcription (see Supplementary Results and Supplementary Figure 6A-C). This variant may also represent a *cis*-element to which *trans*-acting proteins bind to regulate GAA repeat stability^6^. Irrespective of the mechanism, the 5’-CFV may represent an evolutionary protective genomic element insulating the highly mutagenic *FGF14*-SCA27B locus.

Here, we have presented evidence that a sequence variant upstream of the *FGF14*-SCA27B locus is associated with enhanced meiotic stability and thereby appears to offer a protective effect on lineages by reducing the likelihood of expansion. Further study of the *FGF14* flanking variants and identification of additional similar variants across the genome will likely yield further insight into the mechanisms initiating tandem repeat expansion.

## Author contributions

Design or conceptualization of the study: D.P., B.B., S.Z., M.C.D.

Acquisition of data: D.P., G.D.G., M.C., E.D., M.J.D., A.R., V.R., M.W., C.B., C.A., P.J.L., N.G.L., M.R., G.R., H.H., M.S., M.A.E., K.M.B., T.P., B.B., S.Z., M.C.D.

Analysis or interpretation of the data: D.P., G.D.G., M.C., E.D., K.M.B., T.P., B.B., S.Z., M.C.D.

Drafting or revising the manuscript for intellectual content: D.P., G.D.G., M.C., E.D., M.J.D., A.R., V.R., M.W., C.B., C.A., P.J.L., N.G.L., M.R., G.R., H.H., M.S., M.A.E., K.M.B., T.P., B.B., S.Z., M.C.D.

## Acknowledgments

We thank the patients and their families for participating in this study. We thank the Centre d’Expertise et de Services Génome Québec for assistance with Sanger sequencing. We thank Pacific Biosciences Applications lab in Menlo Park, California, USA and Pacific Biosciences bioinformatics team for HiFi sequencing and alignment of the Care4Rare Canada dataset. We also thank Dr. Marek Napierala (University of Texas Southwestern Medical Center, Dallas, TX, USA), Dr. Karen Usdin (National Institutes of Health, Bethesda, MD, USA), Dr. Gabriel Matos-Rodrigues (National Institutes of Health, Bethesda, MD, USA), and Dr. André Nussenzweig (National Institutes of Health, Bethesda, MD, USA) for their helpful comments on the manuscript. D.P. and G.D.G. hold Fellowship awards from the Canadian Institutes of Health Research (CIHR).

## Members of the All of Us Long Reads Working Group

| Member | Affiliation |
| --- | --- |
| Adriana Sedeno Cortes | University of Washington |
| Alan Scott | Johns Hopkins |
| Alba Sanchis-Juan | Broad Institute |
| Allison Rozanski | University of Washington |
| Anant Maheshwari | Broad Institute |
| Andrea Ramirez | National Institutes of Health |
| Andrew Stergachis | University of Washington |
| Anjene Musick | National Institutes of Health |
| Beri Shifaw | Broad Institute |
| Bob Handsaker | Broad Institute |
| Chelsea Berngruber | HudsonAlpha |
| Chris Frazar | University of Washington |
| Chris Lord | Vanderbilt University |
| Chris Rohileck | Broad Institute |
| Christopher Whelan | Broad Institute |
| Colleen Davis | University of Washington |
| David Mohr | Johns Hopkins |
| David Porubsky | University of Washington |
| Dong Ahn Yoo | University of Washington |
| Donna Muzny | Baylor College of Medicine |
| Edibe Nehir Kurtas | Broad Institute |
| Emily LaPlante | Broad Institute |
| Evan Eichler | University of Washington |
| Evie Wan | Broad Institute |
| Fabio Cunial | Broad Institute |
| Fritz Sedlazeck | Baylor College of Medicine |
| Ginger Metcalf | Baylor College of Medicine |
| Glennis Logsdon | University of Washington |
| Harrison Brand | Broad Institute |
| Harsha<br>Doddapaneni | Baylor College of Medicine |
| Hua Ling | Johns Hopkins |
| Hyeonsoo Jeong | University of Washington |
| Isaac Xu | University of Miami |
| James Hwang | Baylor College of Medicine |
| Jane Grimwood | HudsonAlpha |
| Josh Smith | University of Washington |
| Justin Paschall | Johns Hopkins |
| Karynne Patterson | University of Washington |
| Kim Doheny | Johns Hopkins |
| Kiran Garimella | Broad Institute |
| Lee Lichtenstein | Broad Institute |
| Marsha Wheeler | University of Washington |
| Matt Danzi | University of Miami |
| Medhat Madmoud | Baylor College of Medicine |
| Michael Schatz | Johns Hopkins |
| Michael Talkowski | Broad Institute |
| Namrata Gupta | Broad Institute |
| Natthapon<br>Soisangwan | University of Minnesota |
| Niall Lennon | Broad Institute |
| Peng Zhang | Johns Hopkins |
| Philip Empey | University of Pittsburgh |
| PingHsun Hsieh | University of Washington |
| Ryan Lorig-Roach | Broad Institute |
| Sarah Fazal | University of Miami |
| Shadi Zaheri | Broad Institute |
| Shane Neph | University of Washington |
| Shawn Levy | HudsonAlpha |
| Sokny Lim | National Institutes of Health |
| Sophie Schwartz | Broad Institute |
| Stephan Zuchner | University of Miami |
| Steve Huang | Broad Institute |
| Tara Dutka | National Institutes of Health |
| Tristan Shaffer | University of Washington |
| Victoria Popic | Broad Institute |
| William Harvey | University of Washington |
| Winston Timp | Johns Hopkins |
| Xuefang Zhao | Broad Institute |
| Yulia Mostovoy | Broad Institute |

## Online Methods

### Institutional review board approval

The institutional review board of the Montreal Neurological Hospital, Montreal (MPE-CUSM-15-915), the Centre Hospitalier de l’Université de Montréal, Montreal (ND02.045), Clinical Trials Ontario (REB# 1577 CTO), the Children’s Mercy Kansas City (Study #11120514), the Centre Hospitalier Régional Universitaire de Nancy, Nancy (2020PI220), the Center for Neurology, Tübingen (598/2011BO1), and the University of Western Australia, Perth (RA/4/20/1008) approved this study.

### Sanger sequencing

Sanger sequencing of PCR amplification products^7^ was performed at the Centre d’expertise et de services Génome Québec using the ABI3730*xl* DNA Analyzer (Applied Biosystems) and in the Laboratoire de Génétique du Centre Hospitalier Régional Universitaire de Nancy using the ABI3130*xl* DNA Analyzer (Applied Biosystems). Sequences were analyzed using SnapGene v.5.0.8 software (Dotmatics). A total of 339 samples (146 controls and 193 patients with late-onset ataxia) were sequenced. The sequences of 541 of the 678 alleles could be accurately determined and were kept for downstream analysis. Alleles were assessed for the presence of sequence variants in the 5’ and 3’ flanking regions. Motif purity of the *FGF14* GAA repeat locus was assessed using the following ordinal scale: impure (non-GAA motif), low purity (<75% GAA motif), mostly pure (75-99% GAA motif), and pure (100% GAA motif).

### Sizing of expanded *FGF14* alleles

We measured the size of expanded *FGF14* alleles by capillary electrophoresis of FAM-labelled long-range PCR amplification products, as described previously^7^. Amplification products were analyzed on an ABI 3730*xl* DNA Analyzer (Applied Biosystems) with a 50-cm POP-7 capillary using the GeneScan 1200 Liz Dye Size Standard (catalog no. 4379950, Applied Biosystems). Results were analyzed using the GeneMapper software using the built-in microsatellite default settings (version 6.0, Applied Biosystems).

### Targeted nanopore sequencing

Results of Sanger sequencing for samples carrying an *FGF14* expansion were confirmed by means of targeted long-read nanopore sequencing. Nanopore sequencing was performed on 47 individuals with alleles longer than 250 repeat units, as described previously^1^. Among this set, 32 individuals had SCA27B. PCR amplification products were selected for molecular size >400bp using SPRIselect paramagnetic beads for DNA size-selection following manufacturer’s protocol (Beckman Coulter Life Sciences). Pre-sequencing size selection was performed to increase coverage depth of larger alleles. Amplicons were normalized to 150 ng/μl and then multiplexed using native barcoding expansion PCR-free library preparation kits and the SQK-LSK109 sequencing kit as per manufacturer’s instructions, multiplexed and sequenced on the MinION or PromethION platform using the R9.4.1 flow cell (Oxford Nanopore Technologies). Each run included a libprep negative control. Reads were basecalled and demultiplexed with stringent barcodes_both_ends setting using Guppy 5.0.13. Sequences were aligned to the GRCh37 reference human genome using Minimap2-2.17 with the predefined settings for nanopore data. STRique-v0.4.2 was then used to count the number of repeated units observed for each read spanning the *FGF14* tandem repeat site. Motif purity was calculated for each sequencing read as the number of GAA units observed in the portion of the repeat locus-spanning segment of the read divided by the STRique estimation of the total number of repeat units for that read.

### Pacific Biosciences High Fidelity sequencing

The 2,191 samples whose long-read sequencing data was used in this study were drawn from three sources: 1,126 samples spanning 525 families from the Children’s Mercy Research Institute’s Genomic Answers for Kids program, 1,027 samples spanning 1,027 families from the All of Us Long Reads Program Phase 1, and 38 samples spanning 12 families from the Care4Rare Canada research program’s Care4Rare-SOLVE study. The Genomic Answers for Kids dataset included 224 trios, from which 411 meiotic transmission events were able to be confidently resolved. The Genomic Answers for Kids dataset included persons from a wide range of genetic ancestries and largely focused on children with rare diseases and their unaffected family members. The All of Us Long Reads Program Phase 1 dataset was composed of adults unrelated to each other and not known to have a rare disease. All persons in that dataset self-reported as black or African-American. The Care4Rare Canada dataset was mostly of European-descent individuals with various unsolved rare genetic diseases (no patient with SCA27B was included in this dataset) and their relatives, who were often unaffected. Samples from the Genomic Answers for Kids program were sequenced to a coverage of approximately 8-25x using one (most parents) to three (most probands) SMRT cells per sample on a Sequel IIe platform at Children’s Mercy hospital. The All of Us Long Reads Phase 1 Program samples were sequenced to a coverage of approximately 8x using a single SMRT cell per sample on a Sequel IIe platform at the Hudson Alpha Institute. The Care4Rare Canada samples were sequenced to a coverage of approximately 30x for affected individuals (n=26) using three SMRT cells per sample and a minimum of 10x for unaffected family members (n=12) using a single SMRT cell per sample on a Sequel IIe platform at the Pacific Biosciences Applications lab in Menlo Park, California, USA.

All samples were aligned to the GRCh38 build of the human genome. TRGT v0.3.3 or v0.3.4 software^8^ was then run on each sample using default parameters to resolve the sequences of the two alleles in each person. The repeat specification given to TRGT was for the genomic region chr13:102161544-102161756 which includes the *FGF14* GAA repeat locus along with 25-bp of flanking sequence on each side. The alleles called by TRGT were then analyzed for variants in the flanking regions as well as the sequence content of the repetitive region. The repetitive regions were segmented based on fuzzy matching of the repeat motifs, such that up to 1 off-pattern nucleotide was tolerated every 12bp. Then, the GAA purity of each allele was calculated as the proportion of the allele (excluding the flanking sequence) that was spanned by segmentations carrying the GAA motif. The threshold for an allele to be considered GAA-pure was set at 95% purity.

## Data availability

Genomic data from Sanger sequencing have not been consented for sharing. The data created through the All of Us Long Reads Phase 1 Program is available to All of Us Research Program researchers. The Care4Rare-SOLVE data is available through Genomics4RD (https://www.genomics4rd.ca) via controlled access requests. The data created as part of Genomic Answers for Kids is available through NIH/NCBI dbGAP (https://www.ncbi.nlm.nih.gov/projects/gap/cgi-bin/study.cgi?study_id=phs002206.v4). Code for running TRGT and analyzing its output alleles will be made available upon request.

## Web resources

### TRGT software

Dolzhenko, E. PacificBiosciences/trgt: Tandem repeat genotyping and visualization from PacBio HiFi data. Available at: https://github.com/PacificBiosciences/trgt. (Accessed: 21^st^ February 2023)

### RNAStructure Web Server

Bellaousov, S., Reuter, JS., Seetin, MG., Matthews, DH. RNAStructure: web servers for RNA secondary structure prediction and analysis. Available at: http://rna.urmc.rochester.edu/RNAstructure.html (Accessed: 21^st^ February 2023)

