## Supplementary Material for "A common flanking variant is associated with enhanced meiotic stability of the *FGF14*-SCA27B locus"

**Supplementary Information for Pellerin, et al.**

**Supplementary Results**

A total of 4,382 alleles were resolved by PacBio sequencing. Of those, 874 (19.95%) alleles carried the reference 5'-flanking sequence within 3 single nucleotide changes (5'-RFS) and 3,120 (71.20%) alleles carried the 17-bp deletion-insertion common 5'-flanking variant (5'-CFV) (NC_000013.11: g.102161566_102161576delinsTAGTCATAGTACCCCAA). The 5'-flanking sequence in the remaining 388 alleles (8.83%) sequenced by PacBio not carrying the 5'-CFV or the 5'-RFS displayed a range of variations that form another 6 groups. These additional groups mainly consisted of variations in the number of cytosines (Cs) and/or adenine (As) in the final six nucleotides (CCCCAA) of the 5'-CFV. We defined these groups as C1 (NC_000013.11: g.102161566_102161576delinsTAGTCATAGTACAA), C2 (NC_000013.11: g.102161566_102161576delinsTAGTCATAGTACCAA), C3 (NC_000013.11: g.102161566_102161576delinsTAGTCATAGTACCCAA), and C5 (NC_000013.11: g.102161566_102161576delinsTAGTCATAGTACCCCCAA) by the number of cytosines present in that final span (Supplementary Figure 1A). The 17-bp deletion-insertion 5'-CFV corresponds to the C4 group. We observed 30 alleles with the C5 5'-flanking variant (0.68% of 4,382 alleles), 24 alleles with the C3 5'-flanking variant (0.55%), 116 alleles with the C2 5'-flanking variant (2.65%), and 168 alleles with the C1 5'-flanking variant (3.83%). The alleles harboring the C3, C4, or C5 variants generally possessed fewer than 30 GAA repeats (3,159/3,177; 99.43%), while alleles with the C1 or C2 variants all possessed more than 30 GAA repeats (284/284; 100%) (Supplementary Figure 1B). Many of these 5'-flanking sequences also included variants with a single or double terminal adenine. Three C4 alleles not counted as part of the 5'-CFV group were observed with a single terminal adenine. Analyzing the variants by their number of terminal adenines revealed that the only C3 sequence over 30 GAA repeats had a single terminal A (Supplementary Figure 1B). In addition to those groups, 10 alleles (0.23%) had a short 5'-flanking sequence of AACCAACTTTCT$(GAA)_n$ (NC_000013.11: g.102161564_102161576del), all of which carried more than 30 GAA repeats (10/10; 100%). Finally, 37 alleles (0.84%) had a 5'-flanking sequence with other configurations. Together, these data indicate that the 17-bp deletion-insertion 5'-CFV, and any sequence within 1 nucleotide change of it, were overwhelmingly associated with shorter alleles containing fewer than 30 GAA repeats (3,159/3,176; 99.46%). Conversely, 5'-flanking variants with two or more nucleotide changes from the 17-bp deletion-insertion 5'-CFV were associated with sequences longer than 30 repeats (1,183/1,206; 98.09%).

The Sanger data did not contain any occurrences of the C1, C2, C3, or C5 variants observed in the PacBio samples. We observed ten alleles related to the reference 5'-flanking sequence that were more than 3 single nucleotide edits removed from the reference, all of which were longer than 30 repeats.

Since the 5'-flanking region of the *FGF14* (GAA)•(TTC) repeat locus harbored variants related to the repeat length and meiotic stability, we studied the 3'-flanking region for similar patterns. We analyzed the 25 nucleotides immediately following the termination of the $(GAA)_n$ repeat sequence in 2,191 individuals (4,382 alleles) by whole-genome long-read PacBio HiFi sequencing and found that most alleles matched the reference sequence (3,743/4,382; 85.42%). The only common variant observed was an A>C polymorphism (rs61965263) 4 nucleotides after the termination of the GAA repeat sequence: $(GAA)_n$TAGCAA. This variant was observed in 328 alleles (7.49%). No other variant was observed in more than 1% of alleles. All alleles carrying the polymorphism rs61965263 were below 30 repeat units (Supplementary Figure 2). Furthermore, the majority of the alleles with the reference 3'-flanking sequence were also below 30 repeat units. None of the 3'-flanking variants distinguished unique populations of alleles as observed with 5'-flanking variants (Supplementary Figure 2).

A total of 411 meiosis events were measured by PacBio sequencing. Of these, 276 were from parents with the 5'-CFV and 127 were from parents with the 5'-RFS. The eight remaining meiosis events were from cases where the parental alleles contained the less common 5'-flanking sequences: three meioses harbored C1 flanking variants, one harbored a C2 flanking variant, two harbored C5 flanking variants, and two harbored other flanking variants. The 411 meiosis events together show a clear picture of the length-dependent instability of the GAA repeat tract (Supplementary Figure 4A). We observed stable intergenerational transmission in 3 of 4 meiosis events with C1 or C2 flanking variants, 2 of 2 meiosis events with C5 flanking variant, and 1 of 2 meiosis events with other 5'-flanking variants (Supplementary Figure 4B). Analyzing meiotic stability according to motif purity further showed the stabilizing effect of sequence interruption and impurity on transmission of the repeat, particularly when the locus contains over 75 repeat units (Supplementary Figure 5A-F).

We hypothesized that the 5'-CFV could be aiding meiotic stability relative to the reference sequence by preventing the formation of a DNA secondary structure that is more prone to DNA replication and transcription errors. To test this hypothesis, we analyzed the predicted DNA/RNA secondary structure of the 60 nucleotides upstream of the *FGF14* repeat locus in the presence of the 5'-RFS sequence, and the C4 (5'-CFV) and C1 5'-flanking variants using the RNAStructure web server (Supplementary Figure 6A-C). We observed that the 5'-RFS sequence creates an inverted repeat sequence with the 50 bases upstream, which is predicted to form a multi-branched loop. The 5'-CFV sequence appears less likely to form such a structure as a CCCC tract in this sequence creates a point of resistance to this folding since there is no poly-G sequence with which it can pair. This observation provides an explanation for why the 5'-CFV enhances the stability of the *FGF14* repeat locus. In comparison, the C1 variant appears to increase the stability of the secondary structure relative to the C4 (5'-CFV) variant. This observation provides an explanation for why the short C1 or C2 5'-flanking variants are associated with longer repeats.

### Supplementary Figures

#### Supplementary Figure 1

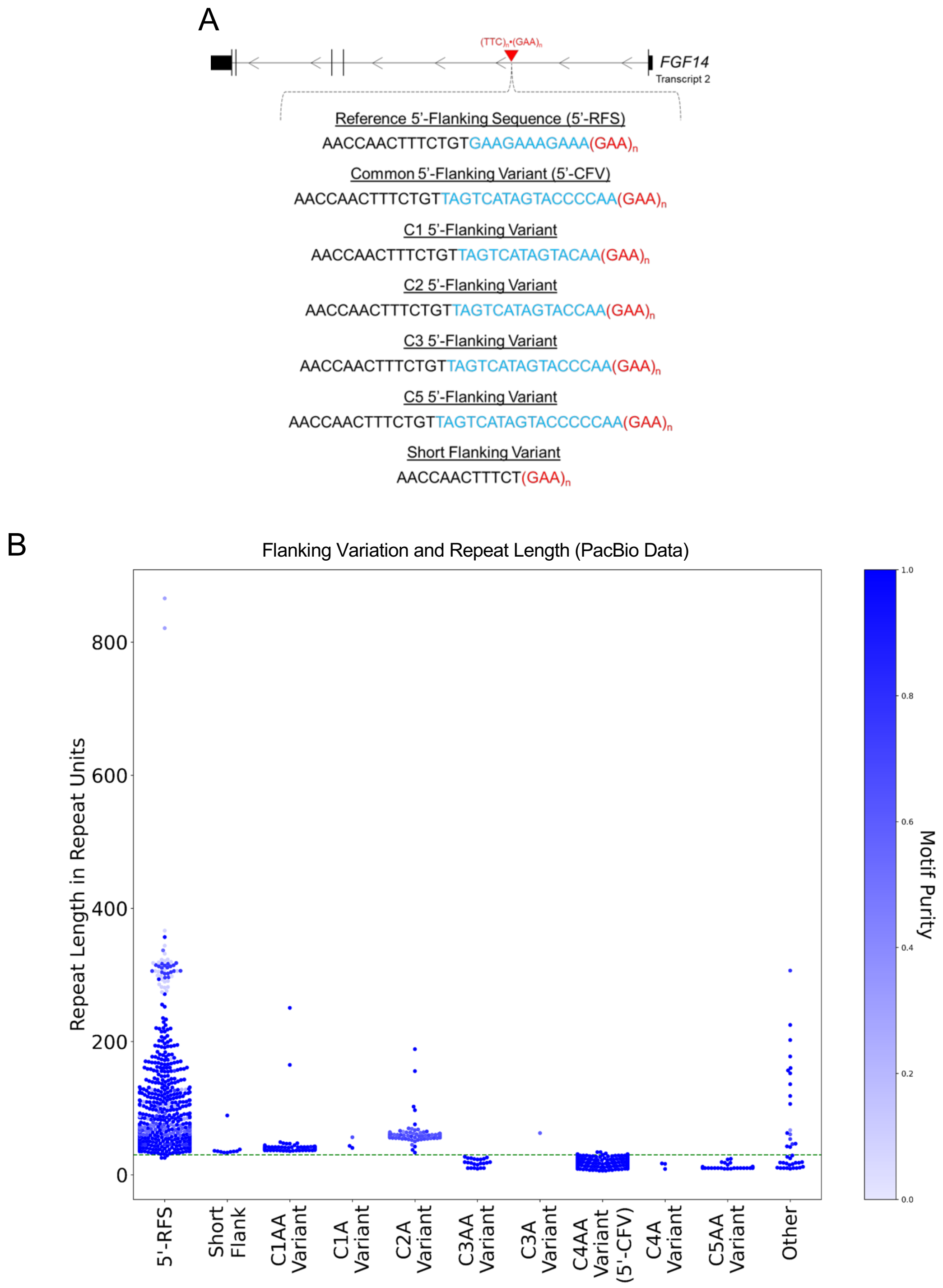


**Supplementary Figure 1:** A) Diagram of the *FGF14* gene with the location of the $(GAA)_n$•$(TTC)_n$ repeat locus in the first intron. The reference 5'-flanking sequence (5'-RFS), the 17-bp deletion-insertion common

5’-flanking variant (5’-CFV; C4 variant), and the C1, C2, C3, C5, and short flanking variant sequences are shown. The 5’-flanking sequence is presented relative to the positive strand for clarity. B) Swarmplot related to Figure 1D showing GAA repeat size as estimated by PacBio HiFi sequencing for 4,382 alleles including each of the C1 through C5 variants, separated into subgroups based on the presence of a single terminal adenine (A) or dual terminal adenines (AA). No alleles with C2AA or C5A 5’-flanking variants were found. This plot also extends the y-axis to show the two alleles that were not plotted in Figure 1D for visual clarity. The color of the data points is a function of the GAA repeat motif purity, with dark blue indicating pure and lighter blue impure motif (a hue scale is shown on the right y axis).

**Supplementary Figure 2**

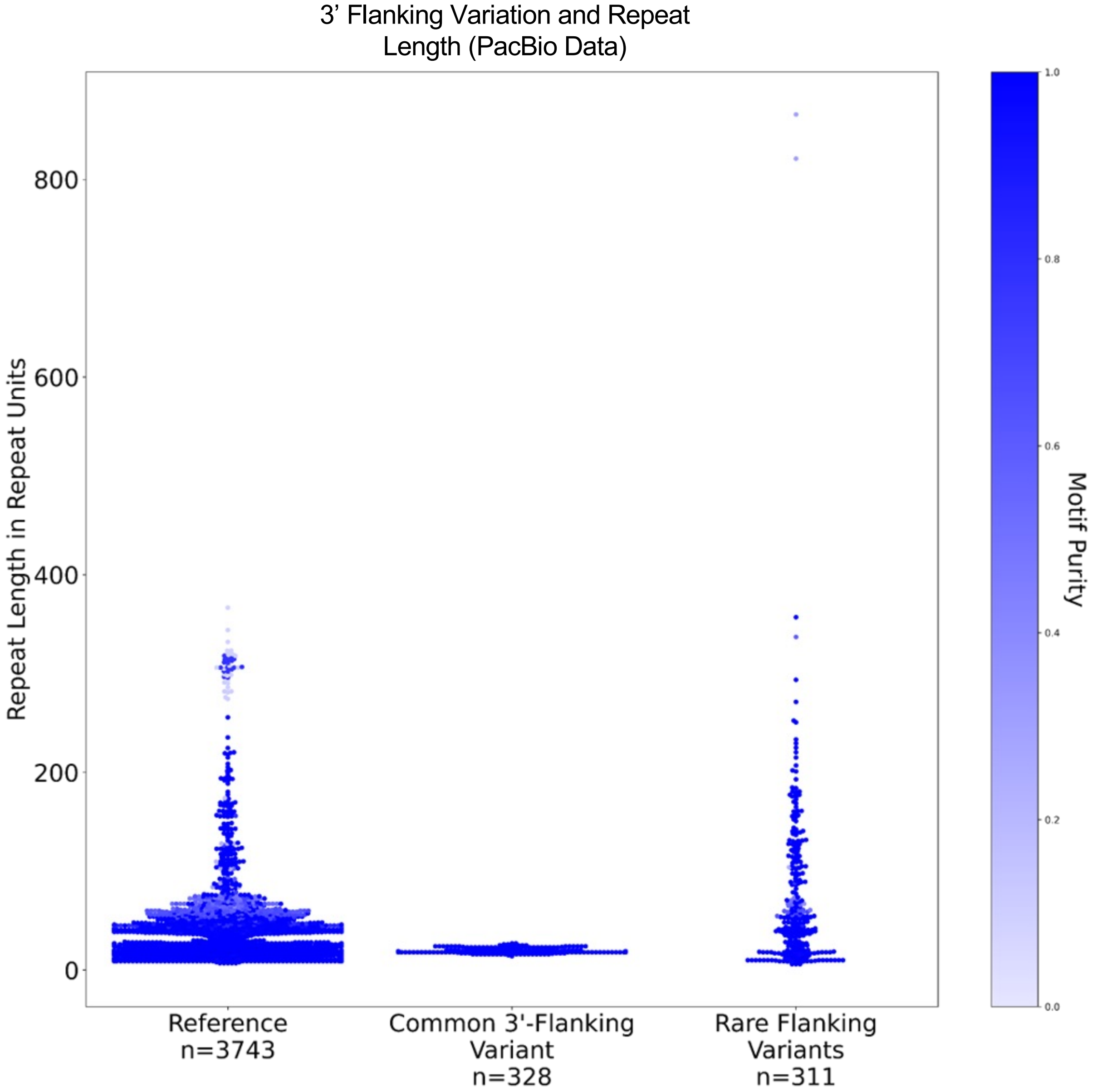


**Supplementary Figure 2:** Swarmplot of the distribution of GAA repeat sizes (in repeat units) for each of the common variations observed in the 3’-flanking sequence of the *FGF14* (GAA)•(TTC) repeat locus. The reference 3’-flanking sequence is $(GAA)_n$TAGAAA, while the common 3’-flanking variant is $(GAA)_n$TAGCAA (rs61965263). All other variants are bundled together into a single category named “rare flanking variants”. The color of the data points is a function of the GAA repeat motif purity, with dark blue indicating pure and lighter blue impure motif (a hue scale is shown on the right y axis).

**Supplementary Figure 3**

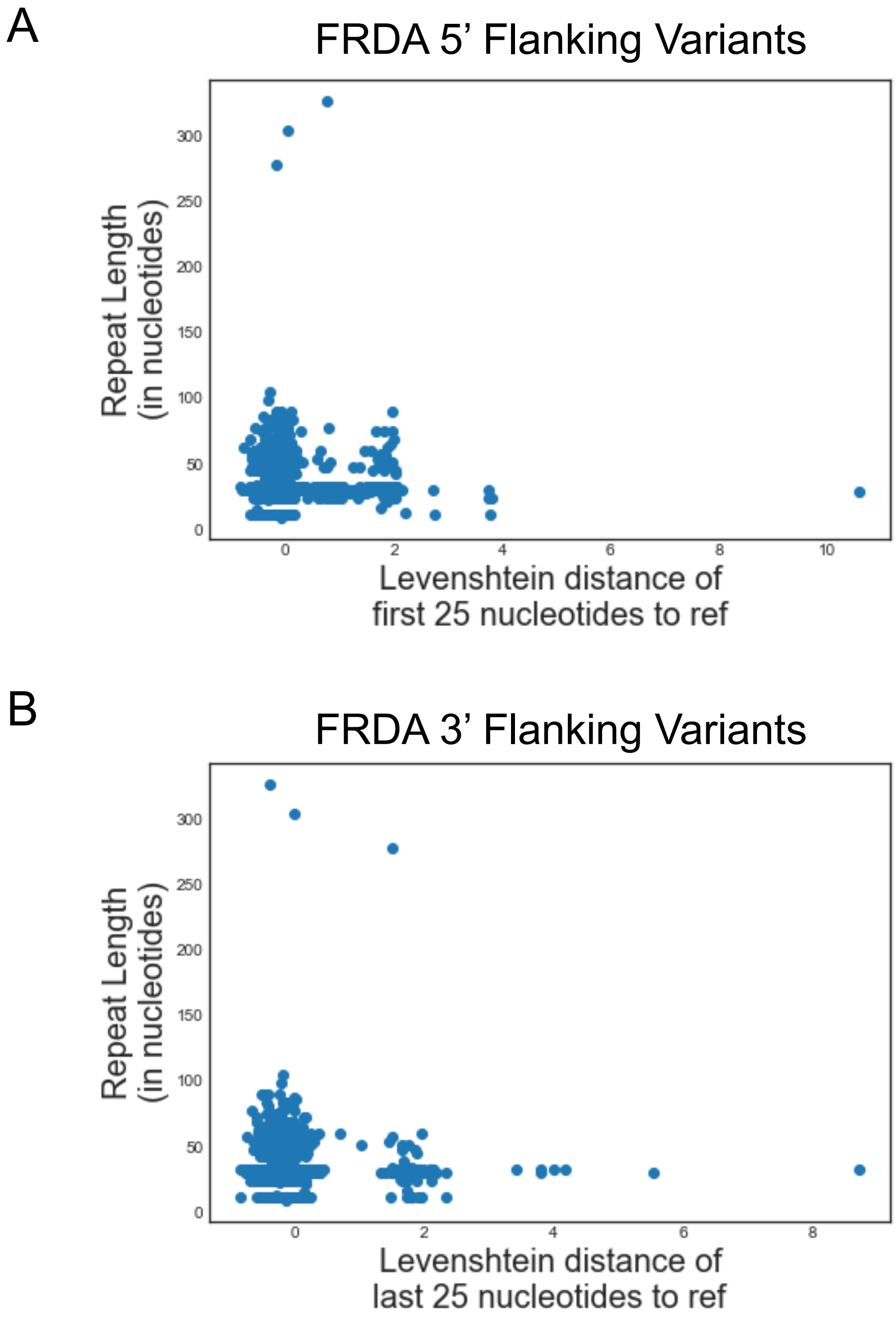


**Supplementary Figure 3:** Scatterplots of 2,054 alleles at the *FRDA* locus where the y-axis plots the GAA repeat length in nucleotides and the x-axis plots the Levenshtein distance between the reference and the observed sequences for the 25 nucleotides immediately A) 5’ to the GAA repeat locus, and B) 3’ to the GAA repeat locus. Gaussian noise was added to the x-axis values to mitigate the extent of dots overlapping each other. No clear segregation of allele sizes by flanking variants was observed.

**Supplementary Figure 4**

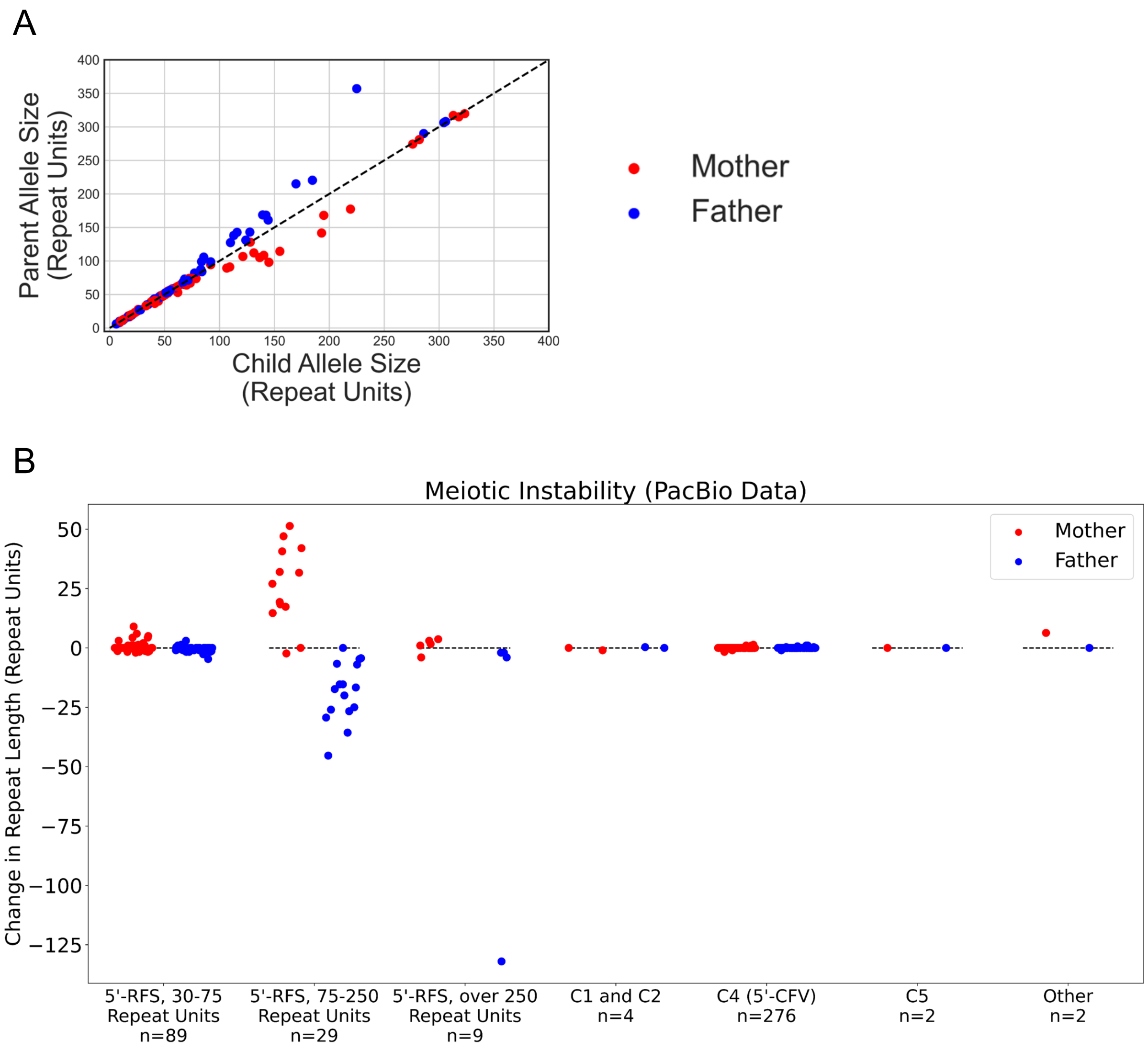


**Supplementary Figure 4:** A) Scatterplot of GAA repeat size as estimated by PacBio for 411 meioses. Contractions are plotted above the dashed identity line while expansions are plotted below that line. B) Strip plot of the change in GAA repeat length across meiosis events as measured by PacBio separated by flanking variant group and parental allele size. Number of meiosis events in each group is stated below the x-axis. The y-axis shows the change in repeat length from parent to child. Contractions are plotted below the dashed lines while expansions are plotted above them. The dotted horizontal lines represent absence of repeat-length variation across meiosis. Random noise was applied across the x-axis within each category to allow visualization of as many data points as possible. This figure extends Figure 2B by plotting the eight additional meiosis events with C1, C2, C3, C5, and other rare 5'-flanking sequences. In panels A and B, red dots are alleles passed from mother to child, while blue dots represent alleles passed from father to child.

### Supplementary Figure 5

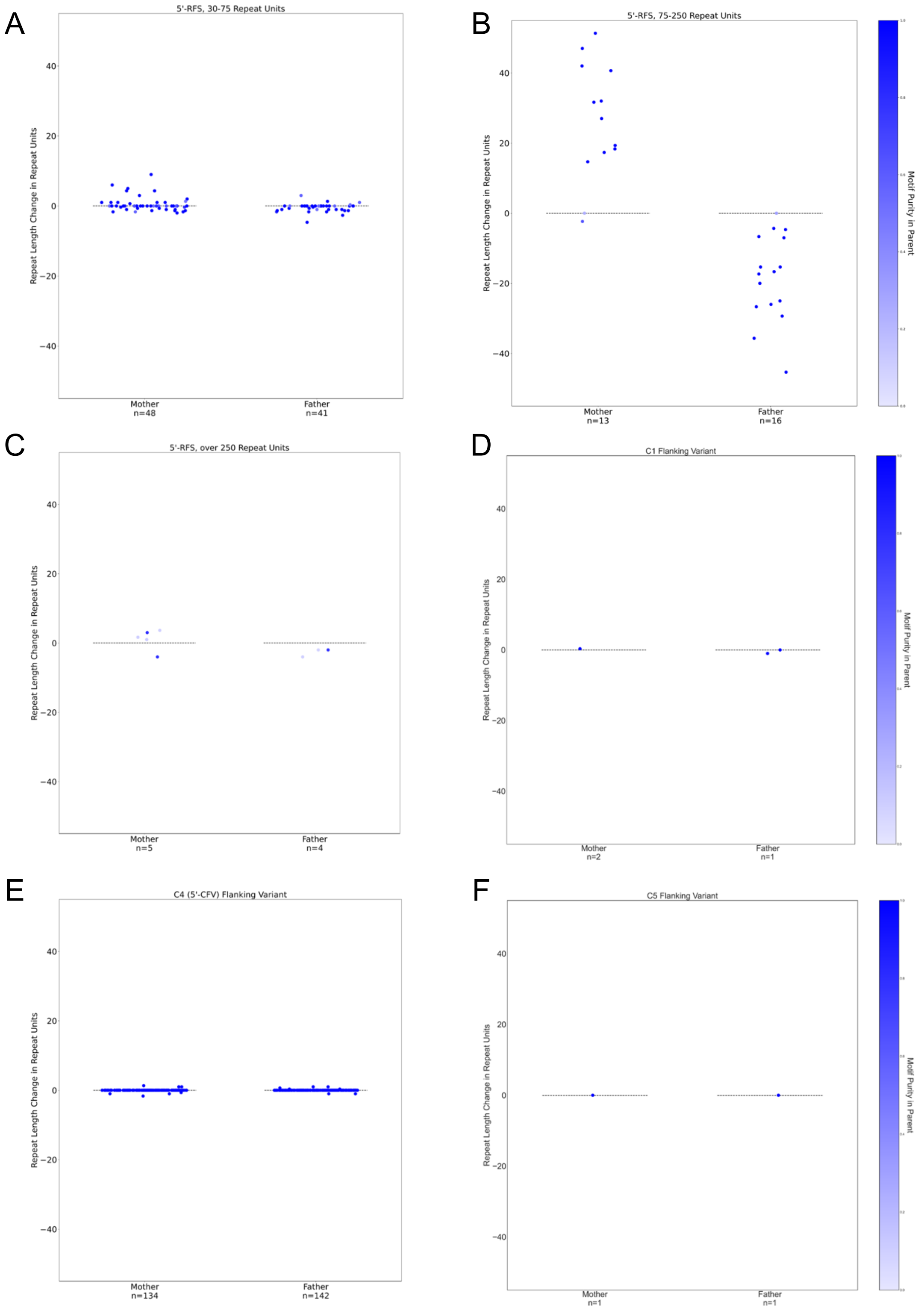


**Supplementary Figure 5:** Strip plots of the change in GAA repeat length across meiosis events. The color of the data points is a function of the GAA repeat motif purity, with dark blue indicating pure and lighter blue impure motif (a hue scale is shown on the right y axis). The y-axis shows the change in repeat length from parent to child. Contractions are plotted below the dashed lines while expansions are plotted above them. The dotted horizontal lines represent absence of repeat-length variation across meiosis. Random noise was applied across the x-axis within each category to allow visualization of as many data points as possible. A) 5'-RFS group of alleles, ranging from 30 to 75 repeat units long. B) 5'-RFS group of alleles, ranging from 75 to 250 repeat units long. C) 5'-RFS group of alleles, over 250 repeat units long. One transmission is not shown here for visual clarity in which a GAA-pure, paternally inherited allele contracted by 132 repeat units. D) C1 group of alleles. E) C4 group of alleles. F) C5 group of alleles. Number of meiosis events in each group is stated below the x-axis.

**Supplementary Figure 6**

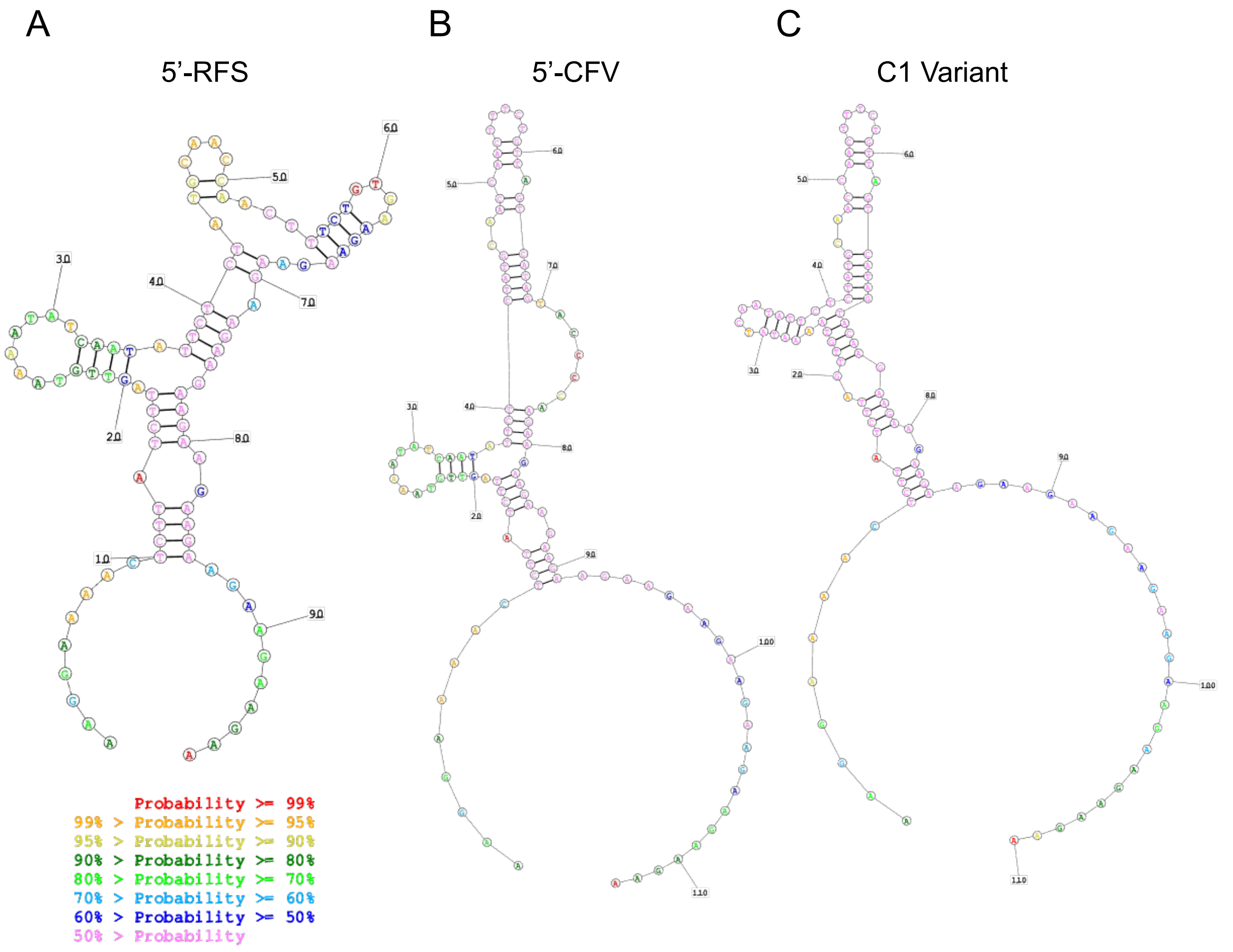


**Supplementary Figure 6:** A-C) Predicted DNA/RNA secondary structure using the RNAStructure web server of the 60 nucleotides upstream of the *FGF14* (GAA)•(TTC) repeat locus along with the first 12 GAA triplet repeats. Panel A shows the most common variant of the reference 5'-flanking sequence, which does not exhibit either of the GAAA sequences immediately prior to the beginning of the GAA repeat. Panel B shows the C4 (5'-CFV) variant. Panel C shows the C1 variant. The probabilities indicate model confidence in correctness of prediction.
